# Identification of a Non-Canonical Ciliate Nuclear Genetic Code Where UAA and UAG Code for Different Amino Acids

**DOI:** 10.1101/2022.12.16.520718

**Authors:** Jamie McGowan, Estelle S. Kilias, Elisabet Alacid, James Lipscombe, Benjamin H. Jenkins, Karim Gharbi, Gemy G. Kaithakottil, Iain C. Macaulay, Seanna McTaggart, Sally D. Warring, Thomas A. Richards, Neil Hall, David Swarbreck

## Abstract

The genetic code is one of the most highly conserved features across life. Only a few lineages have deviated from the “universal” genetic code. Amongst the few variants of the genetic code reported to date, the codons UAA and UAG virtually always have the same translation, suggesting that their evolution is coupled. Here, we report the genome and transcriptome sequencing of a novel ciliate, belonging to the Oligohymenophorea class, where the translation of the UAA and UAG stop codons have changed to specify different amino acids. Genomic and transcriptomic analyses revealed that UAA has been reassigned to encode lysine, while UAG has been reassigned to encode glutamic acid. We identified multiple suppressor tRNA genes with anticodons complementary to the reassigned codons. We show that the retained UGA stop codon is enriched in the 3’UTR immediately downstream of the coding region of genes, suggesting that there is functional drive to maintain tandem stop codons. Using a phylogenomics approach, we reconstructed the ciliate phylogeny and mapped genetic code changes, highlighting the remarkable number of independent genetic code changes within the Ciliophora group of protists. According to our knowledge, this is the first report of a genetic code variant where UAA and UAG encode different amino acids.

## Introduction

The genetic code is one of the most conserved features across life, emerging before the last universal common ancestor [1]. Virtually all organisms use the canonical genetic code which has three stop codons (UAA, UAG, and UGA) and 61 sense codons that specify one of 20 amino acids, including a translation start codon (AUG). Variants of the genetic code, while rare, have been reported in several lineages of bacteria, viruses, and eukaryotic organellar and nuclear genomes [2,3]. Ciliate nuclear genomes are a particular hotspot for genetic code variation. The phylum Ciliophora is a large group of single-celled eukaryotes (protists) that diverged from other Alveolates more than one billion years ago [4]. Ciliates are highly unusual in that they exhibit nuclear dimorphism whereby each cell has two types of nuclei, the germline micronucleus (MIC) and the somatic macronucleus (MAC), each of which contains its own distinct genome structure and function [5]. The MIC genome functions as the germline genome and is exchanged during sexual reproduction. MIC genomes are typically diploid and are transcriptionally inactive during vegetative growth. The MIC genome undergoes rearrangement and excision of micronucleus-limited sequences to serve as a template to generate the transcriptionally active MAC genome [6]. MAC genomes typically contain short, fragmented, gene-dense chromosomes that are present at high ploidy levels (up to tens of thousands of copies) [7].

Known genetic code changes in ciliates involve reassignment of one or more stop codons to specify for amino acids. Most reported ciliate genetic code changes involve reassignment of both the UAA and UAG codons to specify glutamine as in *Tetrahymena*, *Paramecium*, and *Oxytricha* [8], or glutamic acid in *Campanella umbellaria* and *Carchesium polypinum* or tyrosine in *Mesodinium* species [9]. Other known modifications include reassignment of the UGA stop codon to specify tryptophan in *Blepharisma* [8], or cysteine in *Euplotes* [10]. The most extreme example of genetic code remodelling is found in *Condylostoma magnum* where all three UAA, UAG, and UGA stop codons have been reassigned and can specify either an amino acid (glutamine for UAA and UAG, and tryptophan for UGA) or signal translation termination depending on their proximity to the mRNA 3’ end [9,11]. Not all ciliates use non-canonical genetic codes. For example, *Fabrea salina*, *Litonotus pictus*, and *Stentor coeruleus* use the canonical genetic code [9,12].

Changing the meaning of codons from stop to sense requires modifications to the translational apparatus. In eukaryotes, the eukaryotic release factor 1 (eRF1) protein recognises the three standard stop codons in mRNA and triggers translation termination. Studies have shown that mutations in the N-terminus of eRF1 can alter stop codon specificity [8,13–15]. eRF1 specificity to recognise only the UGA codon has evolved independently via different molecular mechanisms at least twice in ciliates with reassigned UAA and UAG codons [14]. Acquisition of tRNA genes with anticodons that recognise canonical stop codons (suppressor tRNAs), via mutations, base modifications or RNA editing enables translation of canonical stop codons into amino acids [16,17].

Tandem stop codons are additional stop codons located in the 3’-UTR within a few positions downstream of a gene in the same reading-frame [18]. They are thought to act as “back-up” stop codons in the event of readthrough, minimising the extent of erroneous protein elongation. For example, in yeast there is a statistical excess of stop codons in the third in-frame codon position downstream of genes with a UAA stop codon [18]. Tandem stop codons have been shown to be overrepresented in ciliates that only use UGA as a stop codon, compared to eukaryotes that use the canonical genetic code [19]. The level of overrepresentation is greater in highly expressed genes [20]. Tandem stop codons are thought to be particularly important in ciliates where, following stop codon reassignment, readthrough events might occur at a higher frequency due to mutations in eRF1 [20].

Several models have been proposed to describe genetic code changes. Under the “codon capture” model, a codon that is rarely used (e.g., due to GC content) is gradually eliminated from the genome followed by loss of the corresponding unused tRNA [21]. Due to random genetic drift the codon could reappear and be captured by a noncognate tRNA charged with a different amino acid, thus changing the genetic code. Such a process would be essentially neutral, not resulting in mistranslated protein products as the codon is eliminated from genes before the change in meaning occurs [17]. Alternatively, under the “ambiguous intermediate” model [22], reassignment of a codon takes place via an intermediate stage, where a codon is ambiguously translated via competing tRNAs charged with different amino acids, or in the context of stop codon reassignment, a suppressor tRNA competing with a release factor. This process would be driven by selection and result in the elimination of the cognate tRNA if the new meaning is advantageous. The “genome streamlining" model is more relevant to small genomes (e.g., organellar genomes or parasites) where there is pressure to minimise translational machinery [23]. More recently the “tRNA loss driven codon reassignment” mechanism was proposed to describe codon reassignments whereby tRNA loss, or alteration of release factor specificity, results in an unassigned codon that can be captured by another tRNA gene [24,25].

In virtually all genetic code changes reported to date, the codons UAA and UAG have the same meaning, i.e., they are either both used as canonical stop codons or are both reassigned to the same amino acid [25]. This suggests that evolutionary or mechanistic constraints couple the meaning of these two codons [26]. One such constraint is wobble binding of a suppressor tRNA gene with a UUA anticodon, where uracil in the first anticodon position can bind to either adenine or guanine in the third codon position of mRNA [27]. Thus, acquiring a suppressor tRNA gene with a UUA anticodon could potentially change the meaning of both the UAA and UAG codons. Wobble binding has been experimentally demonstrated in *Tetrahymena thermophila*, where tRNA-Sup(UUA) was shown to suppress both the UAA and UAG codons, whereas tRNA-Sup(CUA) suppressed only the UAG codon [16]. The first report of nuclear genetic code variants where UAA and UAG have different meanings were reported in transcriptomics analyses where a Rhizarian species (Rhizaria sp. exLh) was shown to use UAG to encode leucine and in a Fornicate (*Iotanema spirale*) where UAG has been reassigned to glutamine [26]. However, in both cases, the UAA codon was retained as a stop codon, thus avoiding the problem of genetic code ambiguity due to wobble binding.

Here, we report the discovery of a novel variant of the genetic code in a ciliate belonging to the Oligohymenophorea class, where the meaning of the UAA and UAG codons have changed to specify different amino acids. Using G&T-Seq [28], we performed parallel genome and transcriptome sequencing of small pools of ciliate cells. Combining genome and transcriptome sequencing data from multiple independently amplified samples enabled co-assembly of a highly complete macronuclear genome assembly and annotation. Genomic and transcriptomic analysis revealed that the UAA codon has been reassigned to specify lysine, while the meaning of the UAG codon has changed to specify glutamic acid. We identified multiple suppressor tRNA genes of both types in the genome, supporting the genetic code changes. We show that UGA codons are significantly enriched in the 3’-UTR of genes suggesting that there is selective pressure to maintain tandem stop codons, which may play a role in minimising erroneous protein elongation in the event of translational readthrough. To our knowledge, this is the first report of a genetic code variant where UAA and UAG specify different amino acids.

## Results & Discussion

### Genome Assembly of an Oligohymenophorean Ciliate

We isolated a novel ciliate species Oligohymenophorea sp. PL0344 from a freshwater pond at Oxford University Parks, Oxford, UK. Attempts to establish a stable long-term culture were unsuccessful so we applied low input single-cell based approaches to generate genomic and transcriptomic data. Small pools of cells (5 – 50 cells) were sorted into a microplate using fluorescence-activated cell sorting (FACS). Parallel genome and transcriptome sequencing was performed using G&T-Seq, which relies on whole genome amplification using multiple displacement amplification (MDA) and transcriptome analysis using a modified Smart-seq2 protocol [28].

A *de novo* genome assembly was generated by co-assembling reads from 10 samples (totalling approximately 6 Gb). Following manual curation and removal of contaminant sequences, the resulting macronuclear genome assembly was 69.7 Mb in length, contained in 3671 scaffolds with an N50 of 59.6 Kb (**Table 1**). Approximately 89% of the corresponding RNA-Seq reads mapped to the genome assembly, indicating high completeness. GC content of the genome is low at 30.6% (**Table 1**), which is similar to previously sequenced ciliate genomes [12]. The mitochondrial genome was also recovered which is a linear molecule 35,635 bp in length with GC content of 25.33% and capped with repeats. The mitochondrial genome contains the small subunit (SSU) and large subunit (LSU) ribosomal RNA (rRNA) genes, 5 tRNA genes, 19 known protein-coding genes, and 13 open reading frames.

**Table 1:**
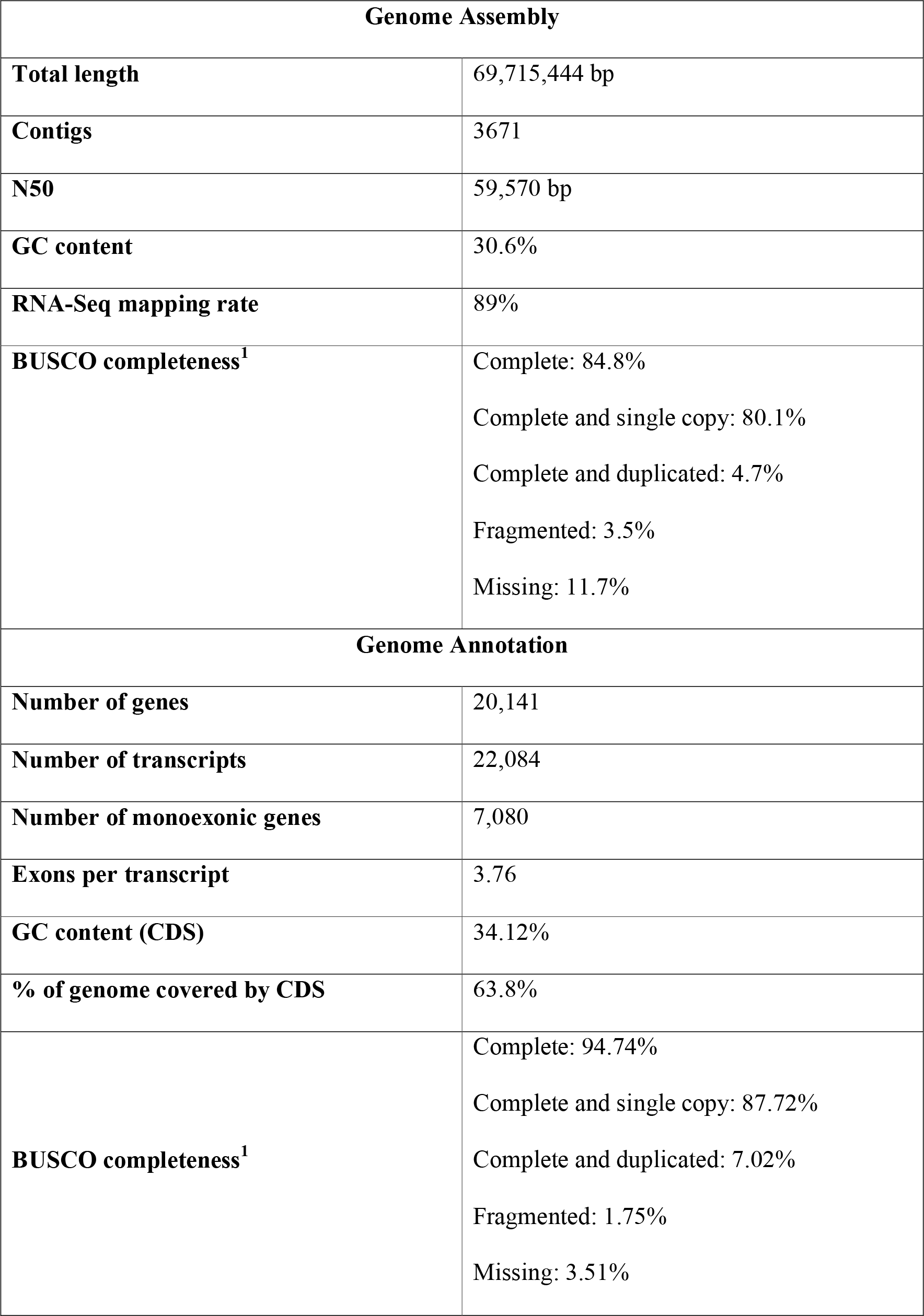
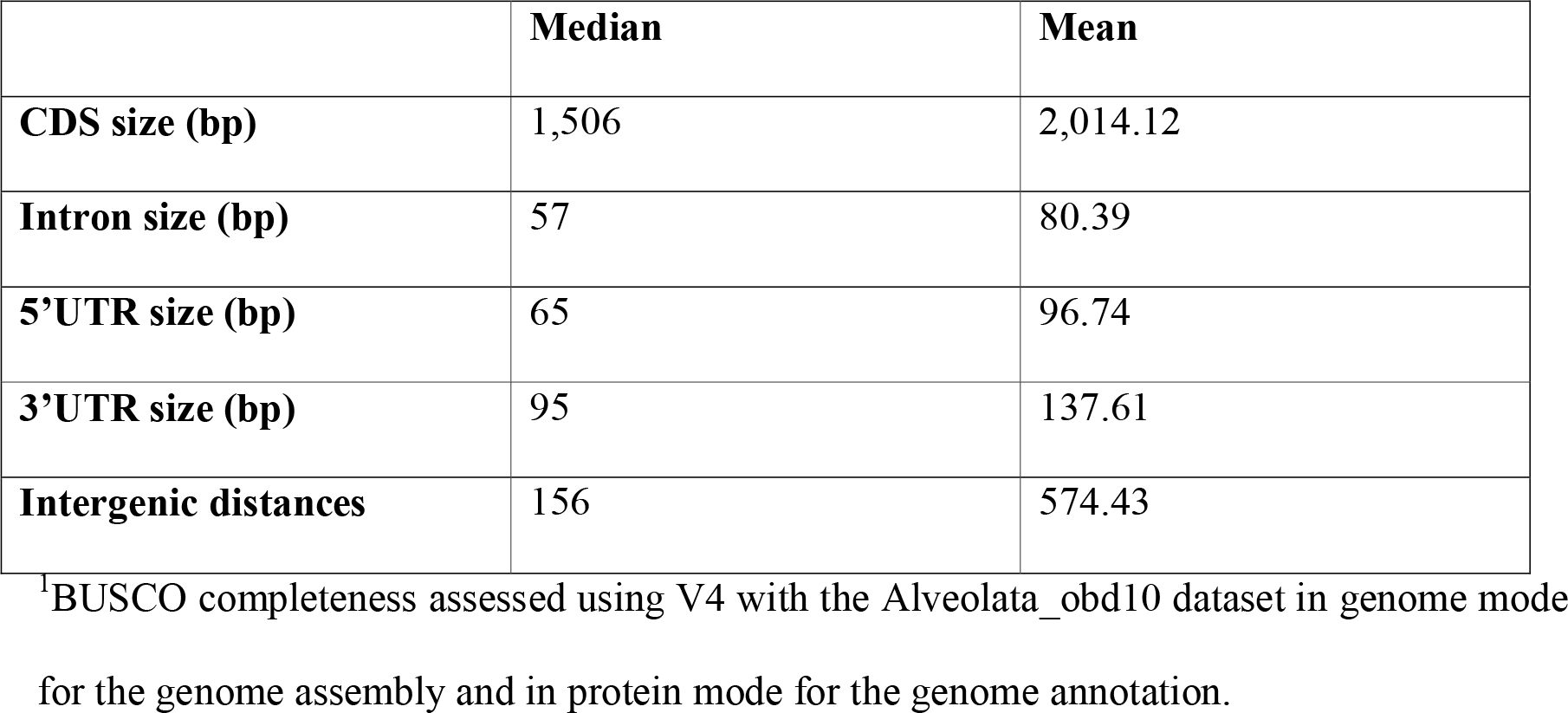
Genome assembly and annotation statistics.

The nuclear encoded SSU rRNA gene sequence is 99.81% identical to an environmental sequence (<u>AY821923</u>) of an unnamed ciliate in the GenBank database, isolated from Orsay, France [29]. Maximum-likelihood phylogenetic analysis of the SSU rRNA gene placed it within a clade containing four unnamed ciliate species (<u>AY821923</u>, <u>HQ219368</u>, <u>LR025746</u>, <u>HQ219418</u>) and *Cinetochilum margaritaceum* (<u>MW405094</u>) with 100% bootstrap support (**Supplementary Figure 1**). Thus, based on the SSU rRNA gene, *C. margaritaceum* is the closest related named species. The SSU rRNA gene of *C. margaritaceum* is 96.03% identical to that of Oligohymenophorea sp. PL0344. *C. margaritaceum* belongs to the Loxocephalida order (Class Oligohymenophorea; Subclass Scuticociliatia), which is considered a controversial order due to its non-monophyly [30,31]. Our phylogenetic analysis places *C. margaritaceum* as a divergent branch relative to other members of Loxocephalida (**Supplementary Figure 1**), which is congruent with previous analyses [30,31], suggesting taxonomic revision is required.

### Oligohymenophorea sp. PL0344 Uses a Novel Genetic Code

Preliminary analysis of the genome sequence revealed that many coding regions contained in-frame UAA and UAG codons. Consistent with codon reassignments in other ciliate species, this suggested that the UAA and UAG stop codons have been reassigned to code for amino acids. Surprisingly however, the meanings of these codons do not match any known genetic code. An example gene (tubulin gamma chain protein), showing six in-frame UAA codons and six in-frame UAG codons, translated and aligned to orthologous protein sequences with representatives from across Eukaryota is displayed in **Figure 1**. Five in-frame UAA codons correspond to highly conserved columns in the alignment where lysine is the consensus amino acid (**Figure 1**). Four in-frame UAG codons correspond to highly conserved columns in the alignment where glutamic acid is the consensus amino acid, and another corresponds to a column with a mix of glutamic acid and aspartic acid (**Figure 1**).

**Figure 1:**
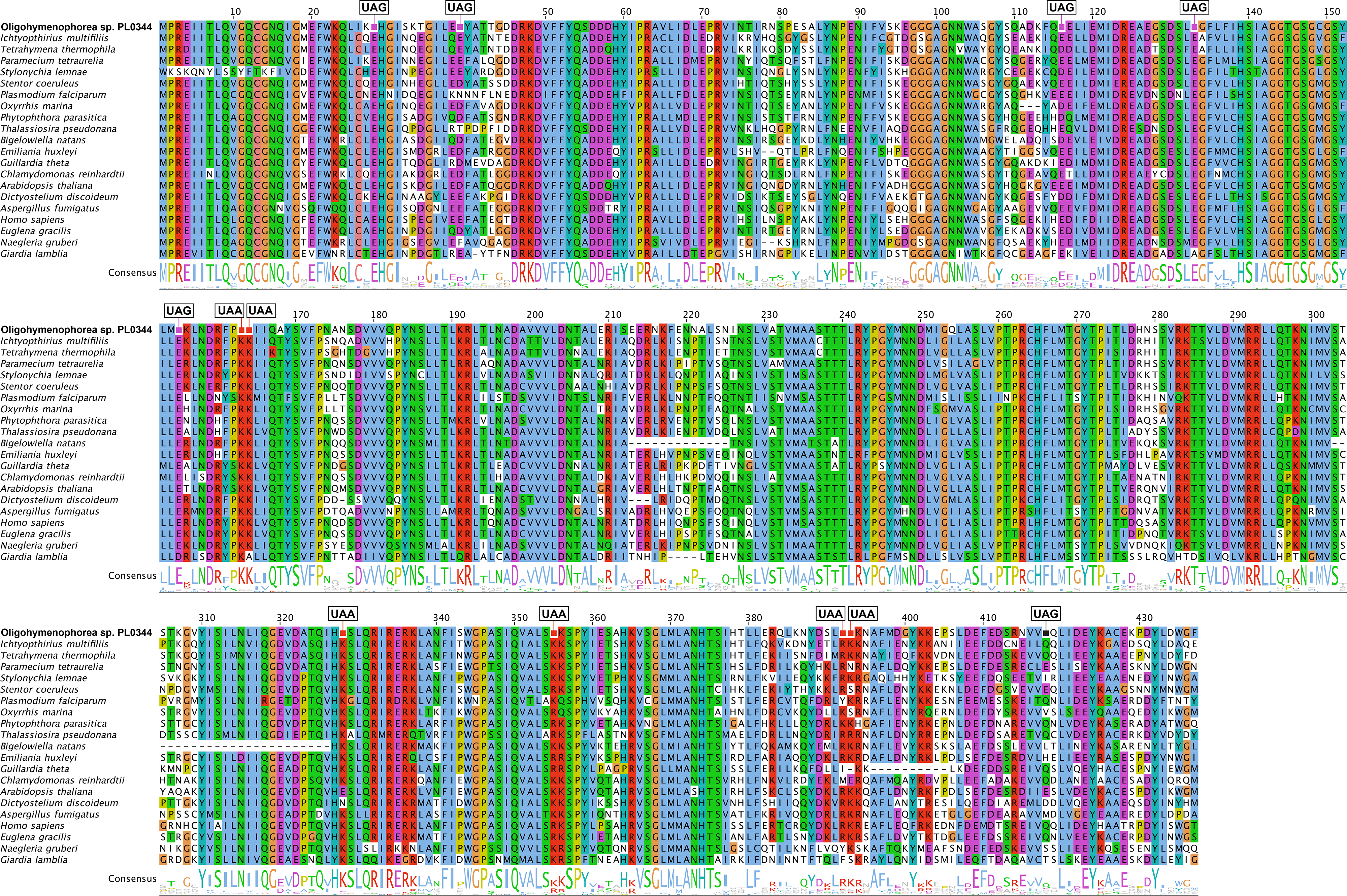
Genetic code change in Oligohymenophorea sp. PL0344. Example multiple sequence alignment of a tubulin gamma chain protein and orthologous sequences spanning Eukaryota. The alignment has been trimmed for visualisation purposes to remove poorly conserved regions and highlight internal UAA and UAG codons.

We used two complementary tools to analyse the genetic code further. First, we used the “genetic_code_examiner” utility from the PhyloFisher package [32], which predicts the genetic code by comparing codon positions in query sequences to highly conserved (> 70% conservation) positions in amino acid alignments from a database of 240 orthologous protein sequences. PhyloFisher identified 58 genes with 87 in-frame UAA codons that correspond to highly conserved amino acid sites. Of these, 74 UAA codons (85%) correspond to highly conserved lysine residues (**Figure 2A**). The second most numerous match was to arginine, another positively charged amino acid, with 9 (10%) hits. PhyloFisher identified 46 genes with 63 in-frame UAG codons that correspond to highly conserved amino acid sites. Of these, 56 UAG codons (89%) correspond to highly conserved glutamic acid residues (**Figure 2B**). The second most numerous match was to aspartic acid, another negatively charged amino acid, with 4 (6%) of hits. Amongst the genes identified by PhyloFisher, 27 contained both in-frame UAA codons and an in-frame UAG codons.

**Figure 2:**
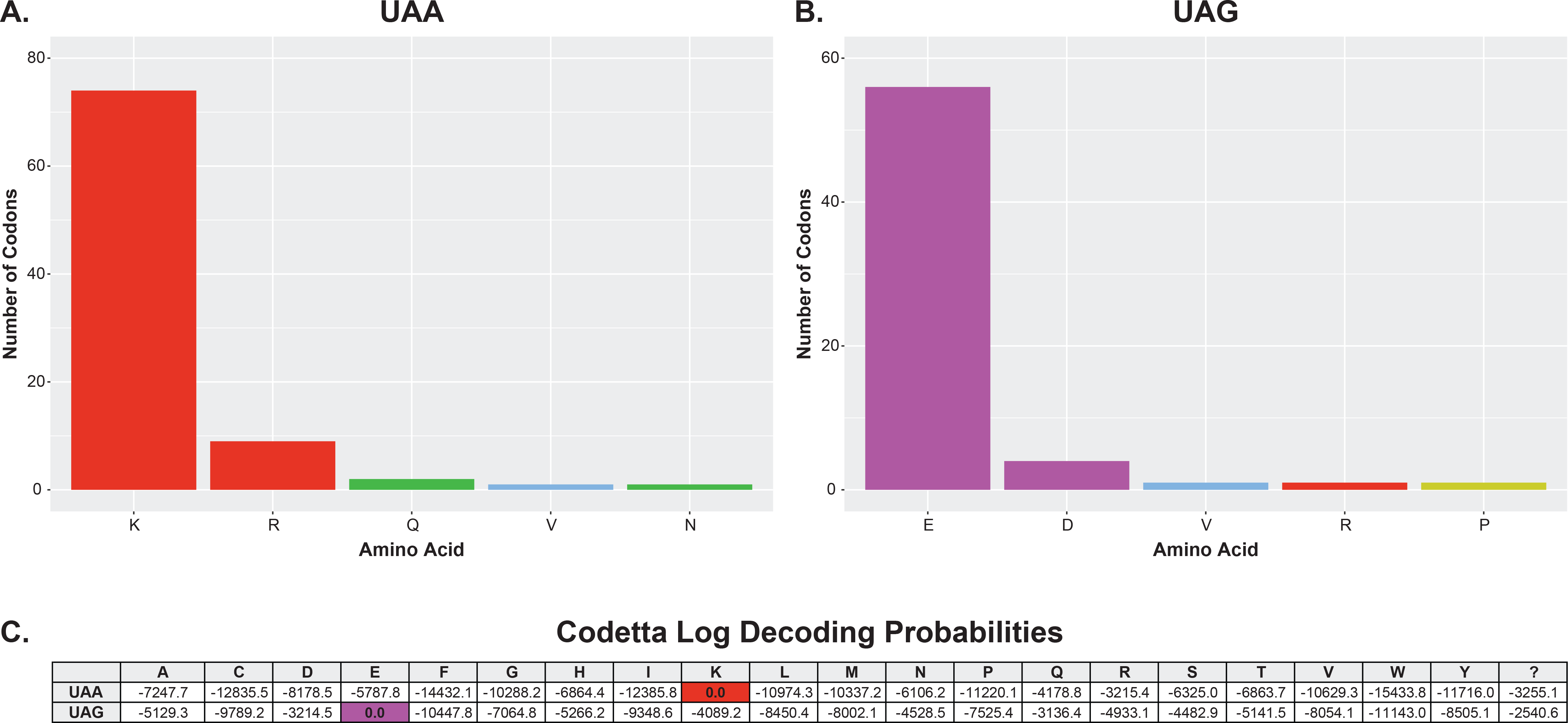
Genetic code prediction for Oligohymenophorea sp. PL0344. PhyloFisher genetic code prediction for the **(A)** UAA and **(B)** UAG codons using the PhyloFisher database of 240 orthologs. Only well conserved (>70%) amino acids are considered. Colours correspond to amino acid properties and match the multiple sequence alignment in Figure 1. **(D)** Codetta genetic code prediction. Log decoding probabilities for the UAA and UAG codons are shown for each of the 20 standard amino acids.

We also analysed the genetic code using Codetta [33,34]. Codetta predicts the genetic code by aligning profile hidden Markov models (HMMs) from the Pfam database against a six-frame translation of the query genome assembly. The meaning of each codon is inferred based on emission probabilities of the aligned HMM columns. From the whole genome sequence, 14,633 UAA codons and 10,160 UAG codons had a Pfam position aligned. Based on these alignments, Codetta also predicted that the UAA codon is translated as lysine and UAG translated as glutamic acid, each with a log decoding probability of zero (**Figure 2C**).

Thus, these results indicate that Oligohymenophorea sp. PL0344 uses a novel genetic code where UAA is translated as lysine and UAG is translated as glutamic acid. This is the first time this genetic code variant has been reported. Furthermore, according to our knowledge, this is the first report of a genetic code variant where UAA and UAG have been reassigned to specify different amino acids. Genetic code variants were previously reported where UAG was reassigned to specify an amino acid (either leucine or glutamine) but UAA was retained as a stop codon in both cases [32]. This is significant as it suggests that the genetic code variant reported herein has overcome mechanistic constraints linking the translation of these two codons.

### Suppressor tRNA Genes

tRNA genes were annotated using tRNAscan [35], resulting in the annotation of 320 tRNA genes, including 15 that are predicted to be pseudogenes. Amongst the annotated tRNA genes are 23 putative suppressor tRNA genes. These are tRNA genes with anticodons complementary to canonical stop codons (UAA, UAG, or UGA). The annotated suppressor tRNA genes include 12 tRNA-Sup(UUA) genes and 10 tRNA-Sup(CUA) genes. tRNAscan also predicted a single tRNA-Sup(UCA) gene, however this was low scoring and was not predicted by ARAGORN [36], an alternative tool to identify tRNA genes. tRNAscan also predicts the function of tRNAs. Many of the tRNAscan isotype predictions were consistent with the predicted genetic code (i.e., UAA = lysine and UAG = glutamic acid), however several putative tRNA genes had low-scoring or inconsistent isotype predictions. To better characterise the suppressor tRNA genes, we compared their sequences to the non-suppressor tRNA genes. Eight of the twelve predicted tRNA-Sup(UUA) genes were most similar to tRNA-Lys genes with UUU or CUU anticodons (68.49% to 80.95% identical) (**Supplementary Table 1**), consistent with the genetic code prediction that UAA has been reassigned to specify lysine. An example tRNA-Sup(UUA) predicted to function as a lysine tRNA is shown in **Figure 3A**. All ten tRNA-Sup(CUA) genes were most similar to tRNA-Glu genes with CUC or UUC anticodons (69.44% to 93.06% identical) (**Supplementary Table 1**), consistent with the genetic code prediction that UAG has been reassigned to specify glutamic acid. An example tRNA-Sup(CUA) predicted to function as a glutamic acid tRNA is shown in **Figure 3B**. Similarly, analysis using phylogenetic networks clusters most of the suppressor tRNA genes with tRNA genes of their predicted function (**Supplementary Figure 2**). We also identified a tRNA gene for selenocysteine, tRNA-SeC(UCA) **(Figure 3C)**, suggesting that the UGA codon is used both as a stop codon and to encode selenocysteine. Thus, all 64 codons can specify amino acids as has been reported in other ciliate genomes [37].

**Figure 3:**
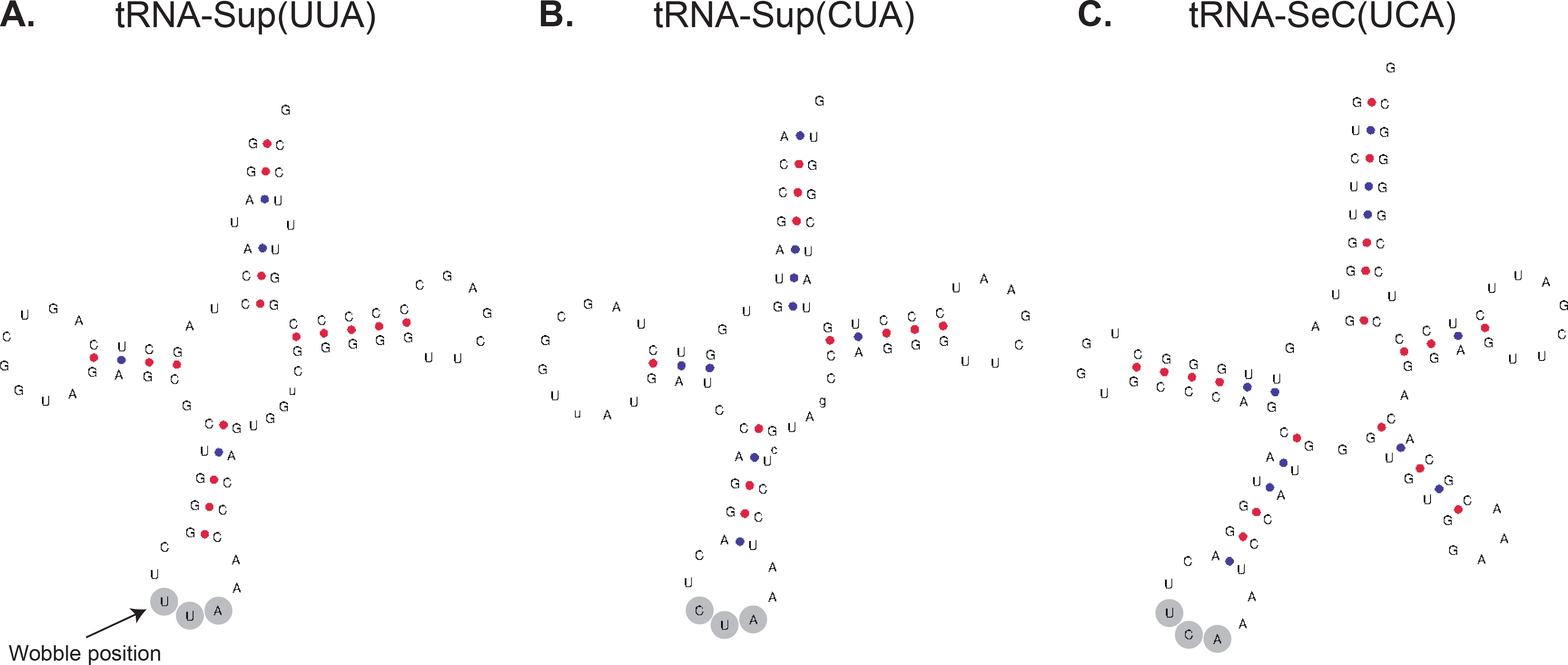
Example tRNA Genes. **(A)** Predicted secondary structure of an example tRNA-Sup(UUA) predicted to function as a lysine tRNA. The wobble position is highlighted. According to wobble-binding rules, uracil at this position can bind to either adenine or guanine in the third codon position of mRNA, allowing the suppressor tRNA to recognise both UAA and UAG stop codons. **(B)** Predicted secondary structure of an example tRNA-Sup(CUA) predicted to function as a glutamic acid tRNA. **(C)** Predicted secondary structure of the tRNA-SeC(UCA) for selenocysteine.

UAA and UAG codons differ only in the wobble position. According to wobble-binding rules, uracil in the first tRNA anticodon position (“wobble position”) (**Figure 3A**) can bind to either adenine or guanine in the third codon position of mRNA [27], allowing tRNA with a UUA anticodon to recognise both UAA and UAG codons. It has been experimentally demonstrated that *T. thermophila* tRNA-Sup(UUA) can recognise both UAA and UAG codons [16]. It has been suggested that wobble binding is a possible explanation as to why UAA and UAG virtually always have the same meaning [9]. Considering that Oligohymenophorea sp. PL0344 has tRNA-Sup(UUA) genes for lysine and tRNA-Sup(CUA) genes for glutamic acid, this raises the question: are UAG codons ambiguously translated as both glutamic acid and lysine? If not, how has it overcome the mechanistic and evolutionary constraints that appear to couple the translation of these two codons? Presumably, if wobble binding allows tRNA-Sup(UUA) to recognise the UAG codon, it would be less efficient than tRNA-Sup(CUA) and outcompeted, possibly resulting in some degree of stochastically translated protein products with glutamic acid residues substituted by lysine at UAG codon positions. Attempts to establish a stable culture were unsuccessful, and while we can overcome this problem to generate a genome assembly using low-input sequencing methods designed for single-cell analysis, such low-input approaches are not available for proteomics. Without proteomics data, it is not possible to determine if UAG is ambiguously translated. Additionally, while the genomic and transcriptomic data provide strong evidence that lysine and glutamic acid are the major translation products of UAA and UAG codons, respectively, we cannot rule out the possibility that other amino acids are (mis)incorporated at these sites which could be detected using mass-spectrometry [38,39]. Furthermore, from suppressor tRNA gene sequences alone, it is not possible to determine if they incorporate modified nucleotides which could alter codon-anticodon binding specificity.

### Genome Annotation and Codon Usage Analysis

Genome annotation incorporating RNA-Seq data and protein alignments from other ciliates resulted in the annotation of 22,048 transcripts from 20,141 gene models (**Table 1**). BUSCO analysis estimates that the genome annotation is highly complete with 94.7% of BUSCO genes recovered as complete, which compares favourably to other high quality ciliate genomes (**Supplementary Table 2**). The median intron size of 57 bp (**Table 1**) is similar to previously sequenced ciliate genomes, such as *Tetrahymena thermophila* and *Oxytricha trifallax* [7,37] but not as short as the extremely short introns (15 – 25 bp) found in *Stentor coeruleus* or *Paramecium tetraurelia* [12,40]. We defined a subset of genes as “highly expressed” based on the 10% of genes with the highest transcripts per million (TPM) values for comparison below. Codon usage is biased towards using codons with lower GC content. This bias is reduced in highly expressed genes which have higher GC content compared to all genes (38.51% versus 34.12%), similar to previous reports in *Paramecium* and *Tetrahymena* [37,41]. The reassigned codons are widely used across genes with 95.9% of genes containing both a UAA codon and a UAG codon. However, their usage is reduced in highly expressed genes (**Supplementary Table 3**). Reduced codon usage in highly expressed genes could indicate translational inefficiency, or that selective pressure to retain canonical lysine and glutamic acid codons is higher in highly expressed genes.

Very little is known about translation termination efficiency in ciliates. This is particularly interesting for ciliates that use only UGA as a stop codon, as UGA is known to be the least robust stop codon and the most prone to translational readthrough [42]. The sequence composition surrounding a stop codon influences the rate of stop codon readthrough. The nucleotide immediately downstream of a stop codon (+4 position) is particularly important, with several studies demonstrating that presence of a cytosine following UGA substantially increases the rate of readthrough [43–45]. Interestingly, examining the sequence composition surrounding stop codons in Oligohymenophorea sp. PL0344, cytosine appears to be avoided following the stop codon (**Supplementary Figure 3**). This is particularly noticeable in highly expressed genes (**Supplementary Figure 3**) where the proportion of genes with a cytosine following UGA is significantly reduced (chi-squared test, p-value = 7.3e-10). This trend has also been observed in *P. tetraurelia* and *T. thermophila* [41]. Tandem stop codons potentially play an important role as “back-up” stop codons, minimising the extent of protein elongation in the event of readthrough [18]. Here, we analysed tandem stop codons by counting UGA codons in the first 20 in-frame codon positions downstream of genes. Our results show that UGA codons are significantly overrepresented (chi-squared test, p-value < 0.05) in the first four in-frame codons downstream of genes (**Figure 4**). 12.3% of genes have at least one UGA codon within the first six in-frame codon positions downstream of genes, similar to the proportion reported for *T. thermophila* (11.5%) where UAA and UAG have also been reassigned to encode amino acids [19]. For comparison, the reassigned UAA and UAG codons are not overrepresented in this region. The frequency of UGA codons at these positions is greater for highly expressed genes whereby 13.6% of highly expressed genes have at least one UGA codon within the first six in-frame codon positions downstream of genes (**Figure 4**). These data add support that there is selective pressure for ciliates with reassigned UAA and UAG codons to maintain tandem UGA stop codons at the beginning of the 3’-UTR. It is tempting to speculate that these additional UGA stop codons play a role in minimising deleterious consequences of readthrough events.

**Figure 4:**
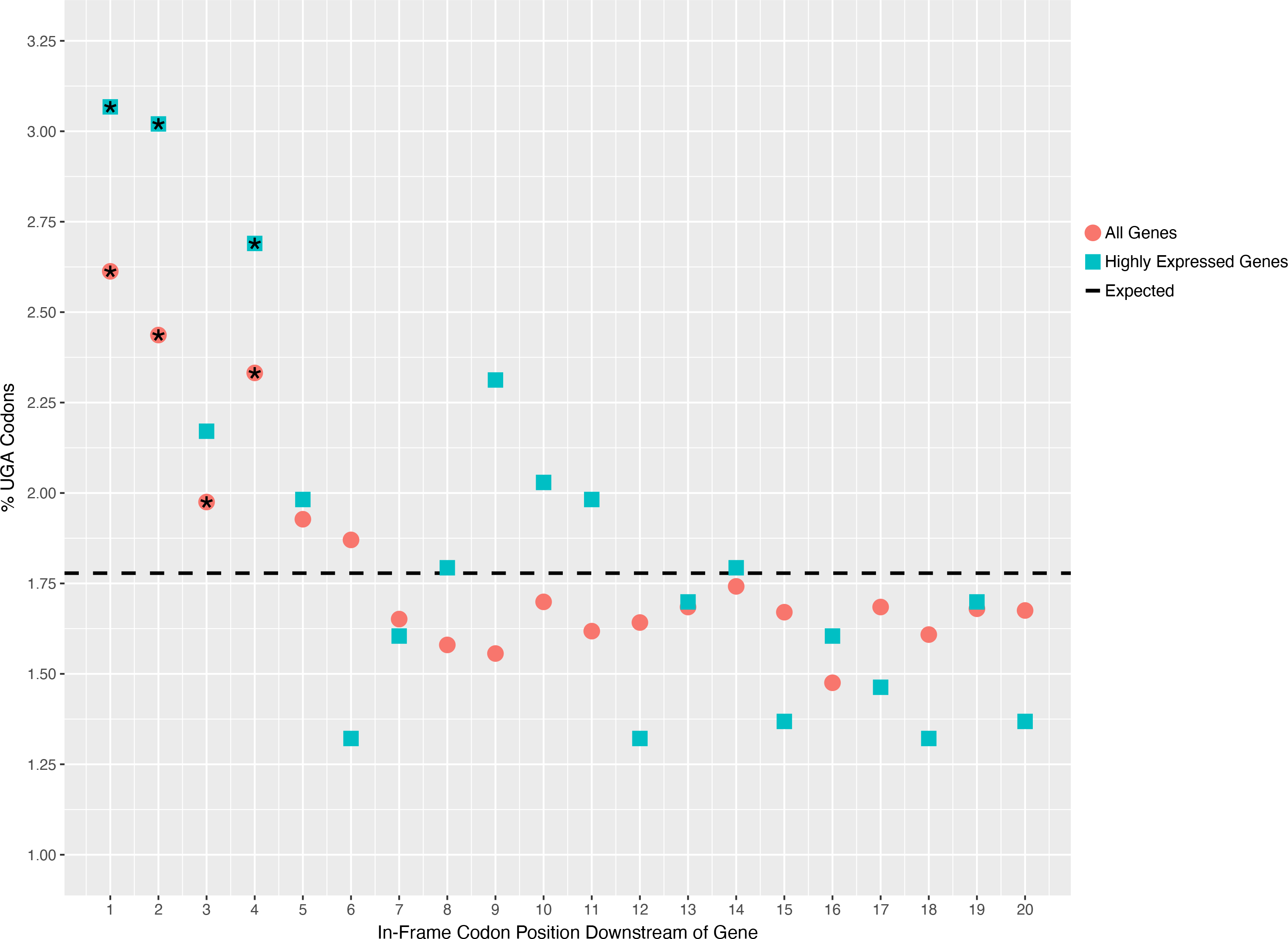
Enrichment of tandem stop codons. The proportion of codon positions occupied by UGA in the 20 in-frame codon positions immediately downstream of all genes and highly expressed genes. Positions where UGA is significantly overrepresented (chi-squared test, p-value < 0.05) are indicated with an asterisk.

### Phylogenomics Analysis of Genetic Code Changes in the Ciliophora

We carried out phylogenomics analyses to map genetic code changes onto the ciliate phylogeny. A phylogenomic dataset consisting of genomic and transcriptomic data from 46 ciliate species and 9 outgroup species was constructed (**Supplementary Table 2**). Phylogenomic reconstruction was performed on a concatenated alignment of 89 single-copy BUSCO proteins (40,289 amino acid sites) using maximum-likelihood (IQ-TREE under LG+F+I+R7 model) and Bayesian (PhyloBayes-MPI under CAT-GTR model) approaches. We also conducted a partitioned analysis on the same dataset using IQ-TREE, with a partitioning scheme which merged the 89 proteins into 14 partitions. The three resulting phylogenies were largely in agreement with each other and with previously published analyses, with full or high support from all three methods at most branches (**Figure 5**). Oligohymenophorea sp. PL0344 was robustly placed within the Oligohymenophorea class in a clade containing Hymenostomatida and *Pseudocohnilembus persalinus* with full support from all three methods (**Figure 5**).

**Figure 5:**
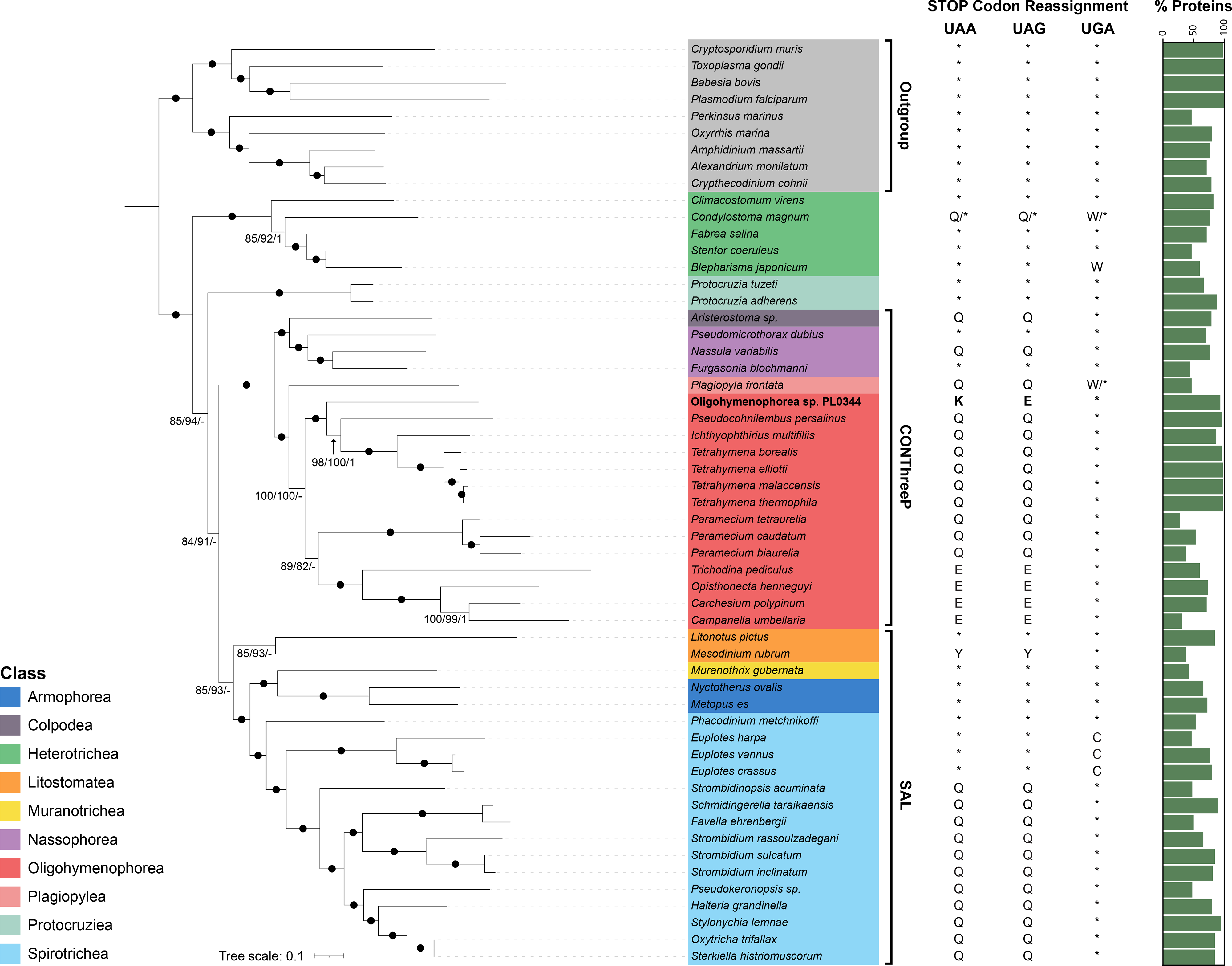
Phylogenomic analysis of genetic code changes in the Ciliophora. Maximum-likelihood phylogeny of 46 ciliate species and 9 outgroup species from the Alveolata, based on a concatenated alignment of 89 BUSCO proteins (40,289 amino acid sites) under the LG+F+I+R7 model using IQ-TREE. The values at branches represent statistical support from 100 non-parametric bootstraps with the LG+F+I+R7 model, 100 non-parametric bootstraps from the IQ-TREE partitioned analysis, and Bayesian posterior probabilities determined under the CAT-GTR model in PhyloBayes-MPI. Branches with full support from all three approaches (i.e., 100/100/1) are indicated with solid black circles. Hyphens indicate branches that weren’t recovered. Stop codon reassignments are shown (*, STOP; Q, glutamine; W, tryptophan; K, lysine; E, glutamic acid; Y, tyrosine; C, cysteine). The percentage of proteins included in the concatenated alignment is shown in the bar plot, highlighting the amount of missing data per species.

The position of *Paramecium* (order Peniculida) is unstable in our phylogenomic analyses. Both the LG+F+I+R7 and partitioned phylogeny group *Paramecium* as sister to the Peritrichia clade with 89% and 82% bootstrap support respectively (**Figure 5**). This is congruent with some previous phylogenomic analyses which recover Peniculida as sister to Peritrichia species [46–48]. However, the Bayesian phylogeny places *Paramecium* as sister to Hymenostomatida and Philasterida (**Supplementary Figure 4**). This grouping has been recovered in some previous phylogenomic analyses [49,50]. The correct placement of Peniculida is unclear based on the current datasets available. The *Paramecium* species included in our analysis have a high proportion of missing data (**Figure 5**). We anticipate that differences in topology may be influenced by varying levels of sensitivity to missing data in the models used. *Mesodinium rubrum* is another problematic taxon which is thought to be prone to long branch attraction (LBA) artefacts. Furthermore, existing *Mesodinium* transcriptomes are contaminated with sequences from their prey [51]. Some previous phylogenomic and phylogenetic analyses place it as an early branching ciliate [52,53], however these may have been influenced by contamination [51]. Here, we account for contamination by removing any sequences from the *M. rubrum* transcriptome with best BLAST hits outside of the Ciliophora (n = 3,574). Both the LG+F+I+R7 and partitioned phylogeny group *M. rubrum* with *Litonotus pictus*, another member of the Litostomatea class, with 85% and 93% bootstrap support respectively (**Figure 5**), while our Bayesian analysis places it as a deep branching ciliate branching before *Protocruzia* (**Supplementary Figure 4**). The grouping of *M. rubrum* with *L. pictus* agrees with a recent phylogenomics analysis of *Mesodinium* species that accounts for LBA and contamination [51].

Where genome or transcriptome assemblies were available, or raw sequencing reads deposited in public databases, we validated the known genetic codes using Codetta and PhyloFisher. All species had the expected genetic code except for *Plagiopyla frontata*. Codetta and PhyloFisher both predicted that UAA and UAG are translated as glutamine in *P. frontata*, which is not surprising given how many ciliate species use this genetic code (**Figure 5**). Interestingly however, both methods predict that UGA has also been reassigned to specify tryptophan in *P. frontata*. From the PhyloFisher dataset of 240 query proteins, 3 (1.25%) contain internal UGA codons that correspond to highly conserved tryptophan residues in other species **(Supplementary Figure 5).** This suggests that *P. frontata* may use UGA both as a stop codon and also rarely as a sense codon to specify tryptophan, similar to the *Condylostoma* genetic code (**Figure 5**) [9,11].

We mapped genetic code reassignments onto the ciliate phylogeny, highlighting the remarkable number of independent genetic code changes within the ciliates (**Figure 5**). Based on our phylogeny, and assuming a non-canonical genetic code doesn’t revert to the canonical genetic code, the translation of UAR (UAA and UAG) codons to glutamine is the most common genetic code variant and has independently evolved at least five times. From our analysis, translation of UGA to tryptophan has independently evolved at least three times in ciliate nuclear genomes. However, it has recently been reported that Karyorelict ciliates (not included in this analysis) use a context-dependent genetic code similar to *Condylostoma*, where UAR has been reassigned to glutamine and UGA specifies either tryptophan or stop depending on context, indicating a fourth independent origin of UGA being translated as tryptophan and a sixth independent origin of UAR being translated to glutamine in ciliates [54]. The translation of UGA to cysteine in *Euplotes*, UAR to tyrosine in *Mesodinium* and UAR to glutamic acid in Peritrichia have all evolved once. The Oligohymenophorea sp. PL0344 genetic code appears to be a relatively recent phenomenon and is unique in that the two codons have different meanings. The Oligohymenophorea class contains at least three different genetic code variants, and no sampled species which have retained usage of UAA or UAG as a stop codon. Our phylogeny suggests that the stop codons UAA and UAG were reassigned to glutamine in the ancestor of Oligohymenophorea (**Figure 5**). These codons were then reassigned to glutamic acid in the Peritrichs, or to lysine (UAA) and glutamic acid (UAG) in Oligohymenophorea sp. PL0344.

It remains unclear why Ciliate genomes are such a hotspot for stop codon reassignments. Our study shows that even within the Oligohymenophorea class, which is relatively well sampled compared to other ciliate clades, there remain novel genetic code reassignments to be discovered. Further sequencing of under-sampled ciliate lineages and other microbial eukaryotes may reveal more variant genetic code changes and help to better understand the evolution and mechanisms of genetic code changes.

## Materials and methods

### Sampling, Ciliate isolation, Culturing, and Cell-sorting

Surface water was collected from a margin of an artificial freshwater pond at Oxford University Parks (51°45’51.0"N 1°15’04.5"W), Oxford (UK) by directly submerging a 1 litre autoclaved glass collection bottle. 200 mL of the water sample were concentrated using a 5 µm filter into a final volume of 20 mL. Oligohymenophorea sp. PL0344 was identified using an inverted microscope (Olympus CKX41) and single cells were isolated manually using a glass micropipette by transferring them into successive drops of 0.2 μm pre-filtered and autoclaved environmental source water. When cells were free of any other eukaryote, they were transferred into a 96 well-plate containing filtered and autoclaved environmental source water. In order to obtain a clonal culture, isolated cells were incubated during a week at 20°C with a 12h:12h light:dark photo-cycle with a photon flux of 32 μmoles·m^-2^·s^-1^. When ciliate cells divided and a dense culture was observed in the well, the mini-culture was scaled-up during a month by successively transferring the cells into larger volumes until a non-axenic but monoeukaryotic ciliate culture of 20 mL of volume was established. Pools of ciliate cells (5 – 50 cells) were then sorted into a 384-well plate containing 5 μL of autoclaved source water using FACS (BD FACSMelody™ Cell Sorter, BD Biosciences). 10 µL of RLT+ lysis buffer (Qiagen) was then added to each well and the plate was sealed and centrifuged (2000 x g, 4 °C, 1 min) to remove bubbles and to ensure that the lysis buffer was at the bottom of each well. The sorted plate was stored at -80°C until processed.

### G&T-Seq, Library Preparation, and Sequencing

Using a magnetic separator, Dynabeads™ MyOne™ Streptavidin C1 (Invitrogen) beads were washed according to the manufacturer’s guidance and then incubated with 2 × Binding &Wash buffer (10 mM Tris-HCl pH 7.5, 1 mM EDTA, 2 M NaCl) and Biotinylated Oligo-dT primer (IDT, 5’-/BiotinTEG/AAG CAG TGG TAT CAA CGC AGA GTA CTT TTT TTT TTT TTT TTT TTT TTT TTT TTT TVN-3’) at 100 µM for 30 minutes at room temperature on a rotator. The oligo-treated beads were washed four times in 1 × Binding &Wash buffer (5 mM Tris-HCl pH 7.5, 0.5 mM EDTA, 1 M NaCl) and then suspended in 1 × SuperScript II First Strand Buffer (Invitrogen) supplemented with SUPERase•In™ RNase Inhibitor (Invitrogen) to a final concentration of 1 U/µl. The lysate was thawed on ice. 10 µl of prepared oligo-dT beads was added to each well containing 12 µl cell lysate. The lysate plate was sealed and incubated on a ThermoMixer C (Eppendorf) at 21 °C for 20 minutes shaking at 1000 rpm. Using a Fluent 480 liquid handling robot (Tecan) and a Magnum FLX magnetic separator (Alpaqua), the lysate supernatant was transferred to a new plate and the beads were washed twice in a custom wash buffer (50 mM Tris-HCl pH 8.3, 75 mM KCl, 3 mM MgCl_2_, 10 mM DTT, 0.5% Tween-20). The supernatant from the washes was added to the left-over cell lysate - containing the genomic DNA - which was stored at -20 °C overnight. The washed beads were suspended in a reverse transcription mastermix of 1 mM dNTPs, 0.01 M MgCl_2_,1 × SuperScript II First Strand Buffer, 1 M Betaine, 5.4 M DTT, 1µM Template-Switching Oligo (5′-AAGCAGTGGTATCAACGCAGAGTACrGrG+G-3′, where “r” prefixes a ribonucleic acid base and “+” prefixes a locked nucleic acid base, Qiagen) then incubated using a ThermoMixer C with the following conditions: 42 °C for 2 minutes at 200 rpm, 42 °C for 60 minutes at 1500 rpm, 50 °C for 30 minutes at 1500 rpm, 60 °C for 10 minutes at 1500 rpm. The cDNA was amplified using HiFi Hotstart Ready Mix (KAPA) and IS Primers to a final concentration of 0.1 µM (IDT, 5’-AAG CAG TGG TAT CAA AGA GT-3’) with the following thermocycling conditions: 98 °C for 3 minutes, then 21 cycles of 98 °C for 15 seconds, 67 °C for 20 seconds, 72 °C for 6 minutes and finally 72 °C for 5 minutes. The cDNA was then purified using 0.8 × vols Ampure XP (Beckman Coulter) and 80% ethanol on the Fluent 480 liquid handling robot and eluted in 10 mM Tris-HCl. The remaining cell lysate was thawed and subjected to a 0.6 × vols Ampure XP clean-up with 80% ethanol. The bead-bound gDNA was isothermally amplified for 3 hours at 30 °C then 10 minutes at 65 °C using a miniaturised (1/5 vols) Repli-g Single-Cell assay (Qiagen). The amplified gDNA was cleaned up with 0.8 × vols Ampure XP and 80 % ethanol, then eluted in 10 mM Tris-HCl. The cDNA and gDNA were quantified by fluorescence (Quant iT HS-DNA, Invitrogen) on an Infinite Pro 200 plate reader (Tecan) then normalised to a final concentration of 0.2 ng/µl in 10 mM Tris-HCl. Dual-indexed sequencing libraries (Nextera XT, Illumina) were prepared using Mosquito and Dragonfly liquid handling instruments (SPT Labtech). The libraries were pooled and cleaned up using 0.8 × vols Ampure XP and 80% ethanol. The libraries were eluted in 10mM Tris-HCl and assessed using a Bioanalyzer HS DNA assay (Agilent), HS DNA Qubit assay (Invitrogen) and finally an Illumina Library Quantification Kit assay (KAPA). Sequencing was conducted on a NovaSeq 6000 with a 300 cycle Reagent kit v1.5 (Illumina) to produce 150 bp paired-end, dual-indexed reads.

### Genome Assembly

Adapter and quality trimming were carried out using BBDuk (https://jgi.doe.gov/data-and-tools/bbtools). Reads which mapped to a database of common lab contaminants (human and mouse) were removed using BBMap. A co-assembly of genomic DNA reads from 10 samples was generated using SPAdes (v3.15.3) [55] with default settings except -k 21, 33, 55, 77 and single-cell mode (--sc) was enabled. The assembly was manually curated and contaminant contigs were removed using a combination of metagenomic binning with MetaBAT2 [56] based on tetra-nucleotide frequencies and taxonomic classification with CAT (v5.2) [57] and Tiara (v1.0.1) [58]. Independent genome assemblies were also generated for all 10 individual samples and average nucleotide identities were compared as a quality control step to confirm that they corresponded to the same species. Assembly statistics were calculated using Quast [59]. Genome completeness was assessed using BUSCO (v4.1.2) [60] with the Alveolata_obd10 dataset run in genome mode.

### Genome Annotation

The genetic code was predicted using Codetta (v2.0) [33,34] and also using the “Genetic Code Examiner” utility from the PhyloFisher package, with the included database of 240 orthologs [32].

Gene models were annotated via the Robust and Extendable eukaryotic Annotation Toolkit (REAT, https://github.com/EI-CoreBioinformatics/reat) and Minos (https://github.com/EI-CoreBioinformatics/minos) using a workflow incorporating repeat identification, RNA-Seq mapping / assembly, alignment of protein sequences from related species and evidence guided gene prediction with AUGUSTUS [61].

A de novo repeat annotation was created using the RepeatModeller [62] v1.0.11 - RepeatMasker v4.07 [63] pipeline with defaults settings and the --gff output option enabled. To ensure high copy number ‘bonafide’ genes were excluded from repeat masking, the RepeatModeler library was hard masked using protein coding genes from 11 ciliate species (detailed below). The protein coding genes were first filtered to remove any genes with descriptions indicating "transposon" or "helicase". TransposonPSI (r08222010) http://transposonpsi.sourceforge.net was then run to remove any transposon hits by hard-masking them and using the filtered gene set to mask the RepeatModeler library. RepeatMasker v4.0.7 was run with the Repbase Alveolata library (RepBaseRepeatMaskerEdition-20170127.tar.gz) and additionally with the filtered RepeatModeler library. The interspersed repeats were combined and used as evidence in the gene build.

The REAT transcriptome workflow was run with RNA-Seq (total 77 million read pairs) from 28 samples. As transcriptome assembly is sensitive to depth of RNA-Seq coverage samples were combined into sets of 28, 10, 10 and 8 samples to ensure reasonable coverage but also allow alternative assemblies to be created. Illumina RNA-seq reads were mapped to the genome with HISAT2 v2.2.1 [64] and high-confidence splice junctions identified by Portcullis [65]. The aligned reads were assembled for each set of samples with StringTie2 v2.1.5 [66] and Scallop v0.10.5 [67]. From the combined set of RNA-Seq assemblies a filtered set of non-redundant gene-models were derived using Mikado [68]. The REAT homology workflow was used to generate gene models based on alignment of proteins from 11 ciliate species (**Supplementary Table 2**). These together with the transcriptome derived models were used to train the AUGUSTUS v3.4.0 gene predictor, with transcript and protein alignments plus repeat annotation provided as hints in evidence guided gene prediction using the REAT prediction workflow. Six alternative AUGUSTUS gene builds were generated using different evidence inputs or weightings for the protein, transcriptome and repeat annotation. The Minos pipeline was run to generate a consolidated set of gene models from the transcriptome, homology, and AUGUSTUS predictions. The pipeline utilises external metrics to assess how well supported each gene model is by available evidence, based on these and intrinsic characteristics of the gene models a final set of models is selected. For each gene model a confidence and biotype classification were determined based on the type and extent of supporting data.

Annotation completeness was assessed using BUSCO (v4.1.2) [60] with the Alveolata_obd10 dataset run in protein mode. tRNA genes were annotated using tRNAscan (v2.0.7) [35]. rRNA genes were annotated using barrnap (v0.9) (https://github.com/tseemann/barrnap).

### Tandem Stop Codon Analysis

To investigate if UGA stop codons are enriched in the 3’-UTR of genes, codon usage of the first 20 in-frame codons downstream of each gene’s stop codon was calculated. Expected frequencies were determined by counting codons in all six reading frames in the 60 bp region downstream of each gene’s stop codon. We also carried out this analysis for highly expressed genes which we defined as the 10% of genes with the highest transcripts per million (TPM) values, calculated using Kallisto [69]. Statistical significance was assessed using the chi-squared test.

### Phylogenetic Analysis of SSU rRNA Genes

Small subunit ribosomal RNA sequences from related species were retrieved from GenBank (**Supplementary Figure 1**). Sequences were aligned using MAFFT (v7.490) with the G-INS-I algorithm [70]. Maximum-likelihood phylogenetic analysis was performed using IQ-TREE (v2.2.0) [71] under the GTR+F+R5 model, which was the best fit model according to ModelFinder [72], with 100 non-parametric bootstrap replicates.

### Phylogenetic Network Analysis of tRNA Genes

A multiple sequence alignment of tRNA genes was generated using MAFFT (v7.490) with the G-INS-I algorithm [70]. tRNA genes predicted to be pseudogenes, containing introns, or that were excessively truncated were excluded. A Neighbour-Net phylogenetic network was constructed using SplitsTree4 [73].

### Phylogenomic Analyses

A phylogenomic dataset of 55 species was assembled including previously published ciliate genomes and transcriptomes with outgroup species from the Alveolata, retrieved from databases and published phylogenomics analyses [74,75] (**Supplementary Table 2**). *De novo* transcriptome assemblies were generated for two species – *Campanella umbellaria* and *Carchesium polypinum*. RNA-Seq reads were retrieved from the sequence read archive (SRR1768423 and SRR1768437) [46]. Transcriptome assemblies were generated using Trinity [76], redundancy was reduced using CD-HIT-EST [77] with an identity cut-off of 98% and protein coding transcripts were predicted using Transdecoder [78]. Coding sequences were translated into amino acids using the correct genetic code (UAR = E). The transcriptome assembly of *Mesodinium rubrum* is contaminated with sequences from its prey. We excluded any *M. rubrum* proteins with a best BLAST hit outside of the Ciliophora to account for this contamination which resulted in the removal of 3,574 (22%) proteins.

BUSCO analysis using the Alveolata_obd10 dataset identified 89 proteins that are present and single copy in at least 65% of species, i.e., at least 36 out of 55 species. Each BUSCO family was individually aligned using MAFFT (v7.490) [70] and then trimmed using trimAl (v1.4) with the “gappyout” parameter [79]. The trimmed alignments were concatenated together resulting in a supermatrix alignment of 40,289 amino acid sites. Maximum-likelihood phylogenetic reconstruction was performed using IQ-TREE (v2.2.0) [71] under the LG+F+I+R7 model, which was the best fitting model according to ModelFinder [72], and 100 non-parametric bootstrap replicates were used to assess branch support. We also conducted a partitioned analysis using IQ-TREE [80] with a partitioning scheme that merged the 89 proteins into 14 partitions with model selection performed by ModelFinder, with 100 non-parametric bootstrap replicates. Bayesian analyses were also performed on the supermatrix alignment using PhyloBayes-MPI (v1.8c) [81] under the CAT-GTR model. Constant sites (n = 3,299) were removed. Two independent Markov chain Monte Carlo (MCMC) chains were run for approximately 12,000 generations. Convergence was assessed using bpcomp and tracecomp with a burn-in of 20%.

## Acknowledgements

This work was funded by Wellcome though the Darwin Tree of Life Discretionary Award (218328) and supported by the Biotechnology and Biological Sciences Research Council (BBSRC), part of UK Research and Innovation, through the Core Capability Grant BB/CCG1720/1 at the Earlham Institute. Part of this work was delivered via the BBSRC National Capability in Genomics and Single Cell Analysis (BBS/E/T/000PR9816) at Earlham Institute by members of the Genomics Pipelines, SingleLCell and Core Bioinformatics Groups, the authors, note the specific contributions of Tom Barker, Vanda Knitlhoffer and Chris Watkins. Additional work was delivered via the BBSRC National Capability in eLInfrastructure (BBS/E/T/000PR9814) at the Earlham Institute by members of the eLInfrastructure group. The authors would like to acknowledge the Scientific Computing group, as well as support for the physical HPC infrastructure and data centre delivered via the NBI Research Computing group. TAR is supported by a Royal Society University Research Fellowship (URF/R/191005).

## Data Availability

All sequencing data and the genome assembly of Oligohymenophorea sp. PL0344 have been deposited to the European Nucleotide Archive under the study accession PRJEB58266. Additional supporting data have been deposited on Zenodo (10.5281/zenodo.7944379).

## Figure Legends

**Supplementary Figure 1:**
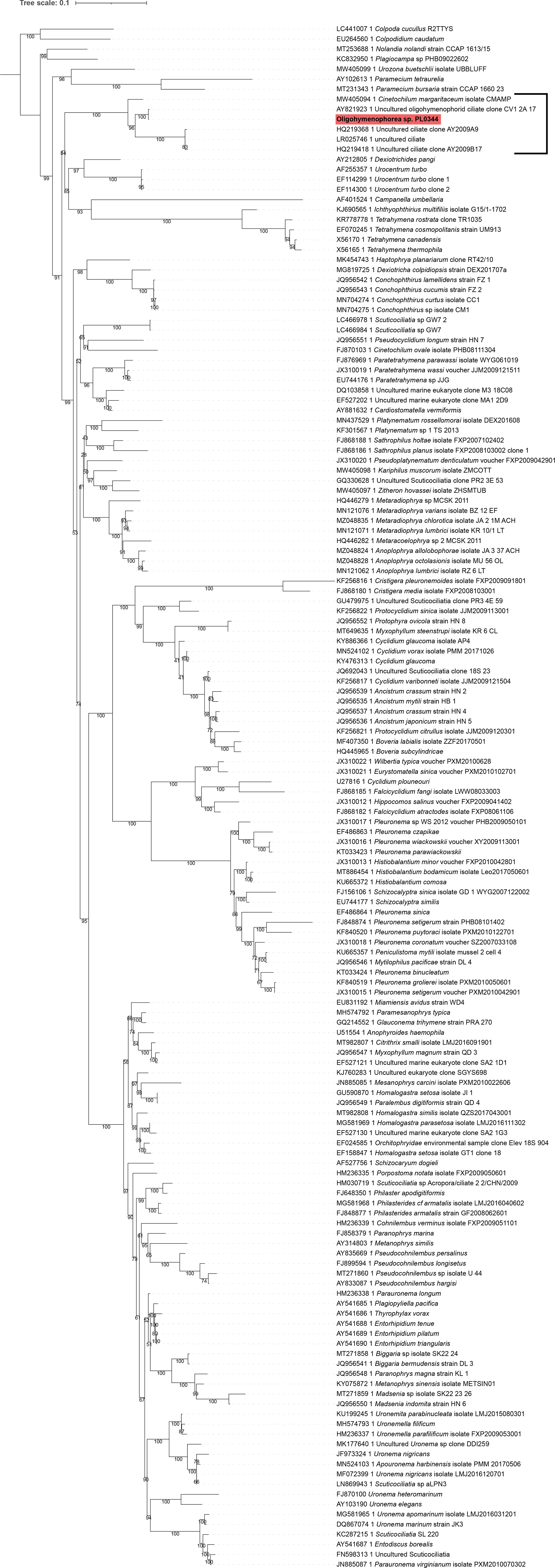
Maximum-likelihood phylogeny of small subunit ribosomal RNA genes under the GTR+F+R5 model using IQ-TREE with 100 non-parametric bootstraps.

**Supplementary Figure 2:**
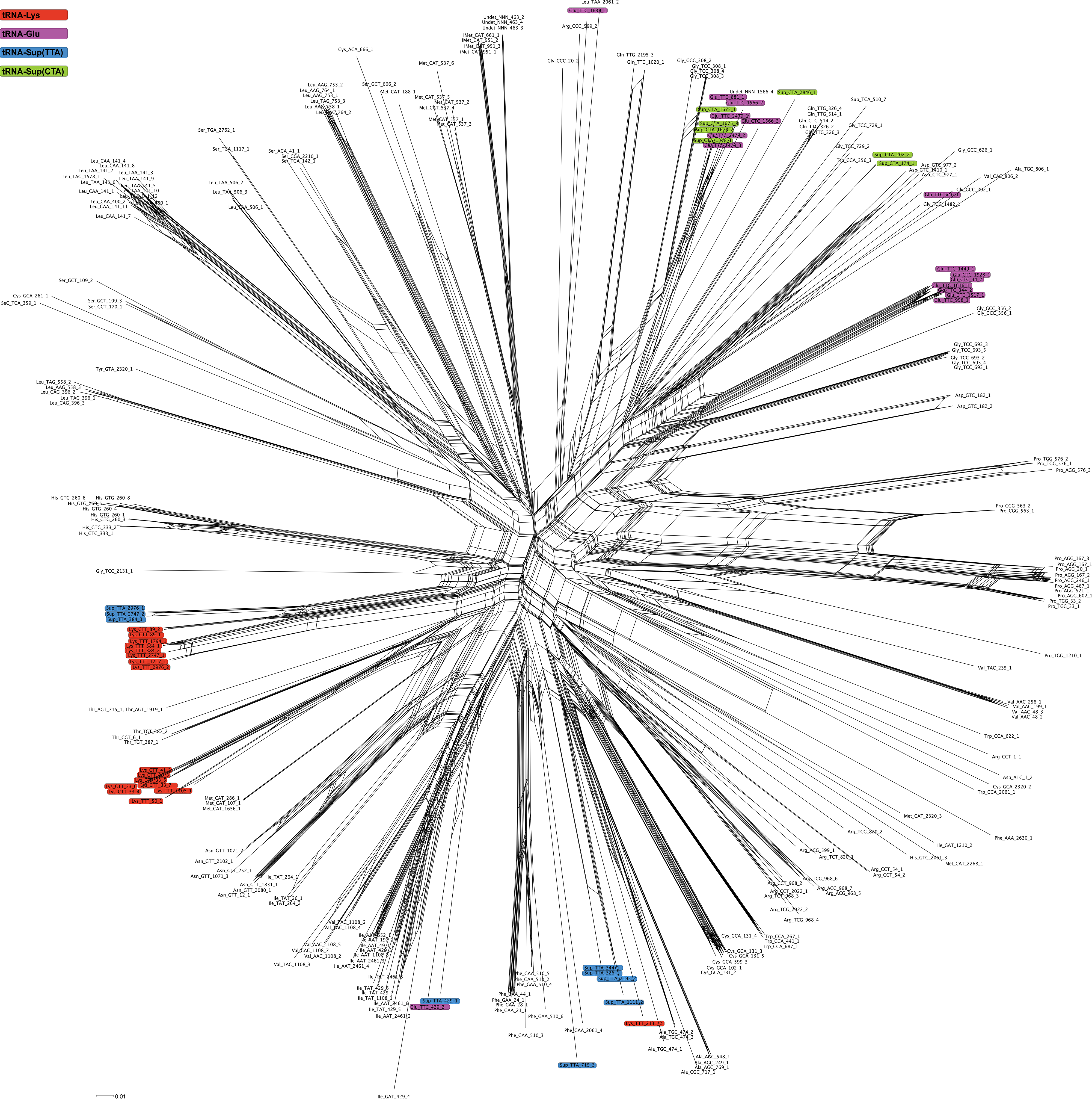
Neighbour-Net phylogenetic network analysis of tRNA genes. tRNA genes predicted to be pseudogenes, contain introns, or that were excessively truncated were excluded. Lysine, glutamic acid, and suppressor tRNA genes are highlighted.

**Supplementary Figure 3:**
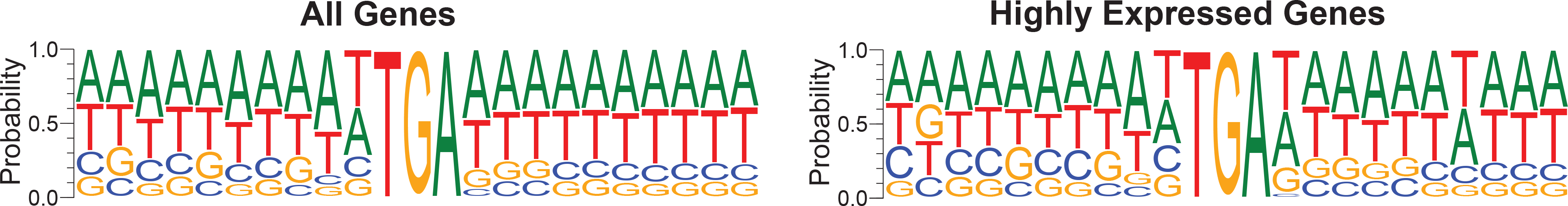
Sequence frequency logo showing the nucleotide composition surrounding stop codons in all genes and in the subset of highly expressed genes.

**Supplementary Figure 4:**
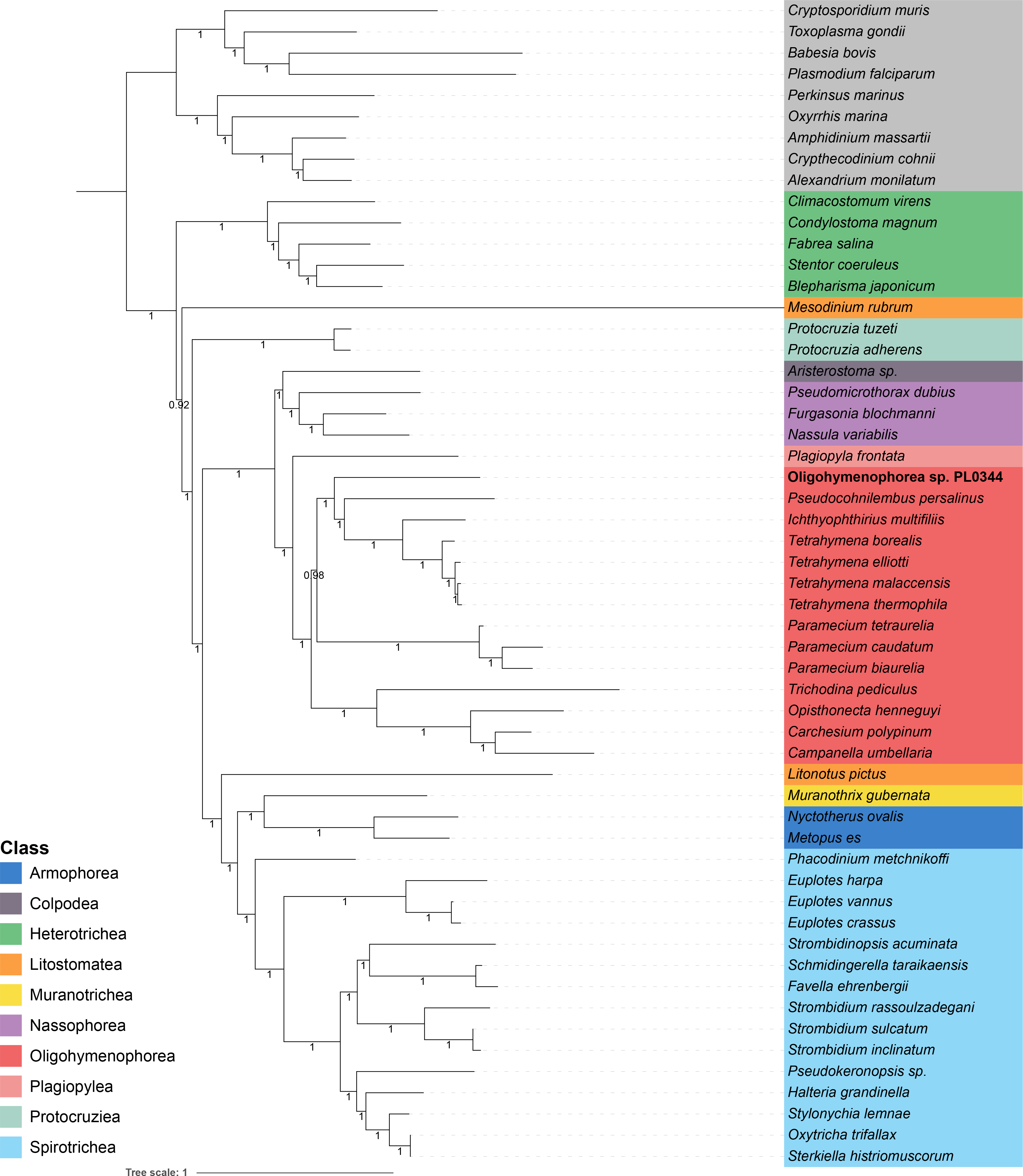
Bayesian phylogenomic analysis of 46 ciliate species and 9 outgroup Alveolata species, based on a concatenated alignment of 89 BUSCO proteins under the CAT-GTR model using PhyloBayes-MPI.

**Supplementary Figure 5:**
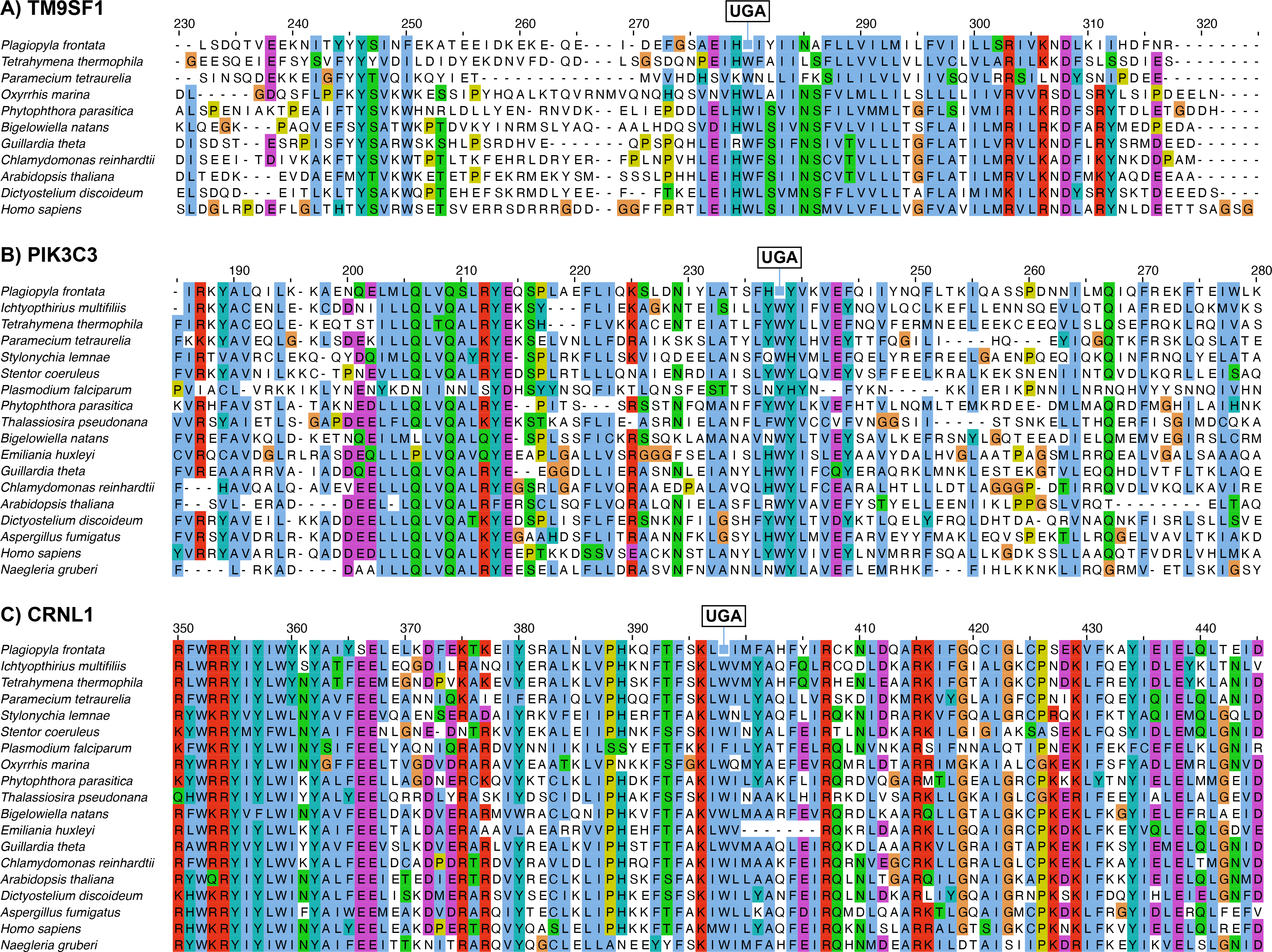
Example multiple sequence alignments of *Plagiopyla frontata* genes with internal UGA codons identified by PhyloFisher with orthologous sequences spanning Eukaryota. **A)** TM9SF1. **B)** PIK3C3. **C)** CRNL1.

## Table Captions

**Supplementary Table 1:** tRNA genes pairwise identities.

**Supplementary Table 2:** Datasets used for phylogenomics and genome annotation.

**Supplementary Table 3:** Amino acid and codon usage.

## Notes

### Competing Interest Statement

The authors have declared no competing interest.

### Summary of Updates

Revised text following peer-review via Review Commons. Added two additional supplementary figures.

